# Disruption of liquid/liquid phase separation in asymmetric GUVs prepared by hemifusion

**DOI:** 10.1101/2024.06.21.600037

**Authors:** Kristen B. Kennison-Cook, Frederick A. Heberle

## Abstract

Model asymmetric bilayers are useful for studying the coupling between lateral and transverse lipid organization. Here, we used calcium-induced hemifusion to create asymmetric giant unilamellar vesicles (aGUVs) for exploring the phase behavior of 16:0-PC/16:1-PC/Cholesterol, a simplified model for the mammalian plasma membrane. Symmetric GUVs (sGUVs) were first prepared using a composition that produced coexisting liquid-disordered and liquid-ordered phases visible by confocal fluorescence microscopy. The sGUVs were then hemifused to a supported lipid bilayer (SLB) composed of uniformly mixed 16:1-PC/Cholesterol. The extent of outer leaflet exchange was quantified in aGUVs in two ways: (1) from the reduction in fluorescence intensity of a lipid probe initially in the sGUV (“probe exit”); or (2) from the gain in intensity of a probe initially in the SLB (“probe entry”). These measurements revealed a large variability in the extent of outer leaflet exchange in aGUVs within a given preparation, and two populations with respect to their phase behavior: a subset of vesicles that remained phase separated, and a second subset that appeared uniformly mixed. Moreover, a correlation between phase behavior and extent of asymmetry was observed, with more strongly asymmetric vesicles having a greater probability of being uniformly mixed. We also observed substantial overlap between these populations, an indication that the uncertainty in measured exchange fraction is high. We developed models to determine the position of the phase boundary (i.e., the fraction of outer leaflet exchange above which domain formation is suppressed) and found that the phase boundaries determined separately from probe-entry and probe-exit data are in good agreement. Our models also provide improved estimates of the compositional uncertainty of individual aGUVs. We discuss several potential sources of uncertainty in the determination of lipid exchange from fluorescence measurements.

**Statement of Significance:** We used calcium-induced hemifusion to create an asymmetric lipid distribution in giant unilamellar vesicles that are models for the mammalian plasma membrane. Confocal fluorescence micrographs of asymmetric vesicles showed that coexisting liquid-ordered and liquid-disordered domains initially present in symmetric vesicles were disrupted after 75% of the saturated lipid in their outer leaflets was replaced with unsaturated lipid. We developed quantitative models for extracting valuable information from the data, including the location of the phase boundary and the compositional uncertainty of individual asymmetric vesicles. The methodology we describe can help reveal the molecular determinants of interleaflet coupling of phase behavior and thus contribute to a better understanding of lipid raft phenomena.

## 1. Introduction

The plasma membrane (PM) separates the cell interior from the outside environment and is centrally involved in many biological processes. In eukaryotic cells, the inner (cytosolic) and outer (exoplasmic) PM leaflets are compositionally asymmetric: the outer leaflet is a mixture of unsaturated phosphatidylcholine (PC) and saturated sphingomyelin (SM) lipids, while the inner leaflet is a mixture of PC, phosphatidylserine, phosphatidylethanolamine, and phosphatidylinositol lipids, nearly all of which are unsaturated^1^. The PM of mammalian cells also contains 30-50 mol% cholesterol^2^ which, unlike phospholipids, can rapidly transit between leaflets^3–6^. While the transbilayer distribution of cholesterol has proven difficult to determine^7^, recent measurements indicate that the majority resides in the outer leaflet^8, 9^.

The striking compositional difference between the inner and outer leaflets raises important questions about lipid miscibility in the PM. Diverse in vitro and in vivo evidence suggests that the outer leaflet, having substantial amounts of both ordered and disordered lipids in addition to cholesterol, may be primed for liquid-liquid phase separation^10^. The propensity of outer leaflet lipids to demix dovetails with the membrane raft hypothesis, which proposes a key biological role for dynamic, nanoscale lipid clustering in processes such as signal transduction, membrane trafficking, and viral entry and exit^11,12^. In contrast to the outer leaflet, most inner leaflet lipids are unsaturated and fully fluid at physiological temperatures, showing little or no tendency toward phase separation in model membrane studies^13, 14^. A key question is whether microdomain formation in the PM outer leaflet can induce segregation of inner leaflet lipids and thereby orchestrate the spatial organization of cytosolic proteins that are anchored to the inner leaflet via lipidation^15, 16^. Equally intriguing is the possibility that the inner leaflet may suppress domain formation in the outer leaflet which, like a loaded spring, would serve as a store of potential energy in the form of unfavorable nearest neighbor interactions. In this scenario, subtle changes in the composition of either leaflet (for example, induced by the activation of scramblases) could act like a switch that triggers lipid and protein reorganization in both leaflets^9^. Investigating these possibilities requires robust methods for preparing asymmetric bilayers where the composition of each leaflet can be precisely controlled^17^.

To this end, several techniques for preparing asymmetric vesicles in a range of sizes have been described in the literature (recently reviewed in Krompers and Heerklotz^18^ and St. Clair et al.^19^). Methods for preparing asymmetric giant unilamellar vesicles (aGUVs) are particularly valuable for optical microscopy studies of phase behavior and include variations of cyclodextrin-mediated lipid exchange,^20,21^ water-oil phase transfer,^22^ and hemifusion.^23, 24^ The latter of these uses millimolar calcium concentrations to induce hemifusion between symmetric GUVs (sGUVs) and a supported lipid bilayer (SLB), thus facilitating exchange of their outer leaflet lipids (Ca^2+^ is removed by EDTA in the final step). Different fluorescent lipid probes incorporated in the sGUV and SLB are used both to visualize the resulting aGUVs and to quantify the extent of outer leaflet exchange. The hemifusion procedure offers some advantages compared to other aGUV preparation methods. Most importantly, it avoids the use of cyclodextrins, organic solvents, or oil that can potentially interact with the bilayer and alter its properties^25^.

While the number of studies utilizing calcium-induced hemifusion is still small^23, 26–29^, it is notable that all published data show wide variability in lipid asymmetry within a population of aGUVs, ranging from very little to nearly complete outer leaflet exchange in a single preparation. As a serendipitous consequence, the degree of asymmetry is a “built-in” variable that can lead to new insights. For example, a recent study from our group investigated the behavior of aGUVs formed by the hemifusion of phase separated DPPC/DOPC GUVs to an SLB composed of DOPC^28^. Sorting the aGUVs in order of increasing outer leaflet exchange revealed a phase transition at approximately 60% exchange: beyond this threshold, most aGUVs were visually uniform in fluorescence images, suggesting an abrupt lateral lipid reorganization arising from interactions between the leaflets. The same study estimated a relative uncertainty of ± 12-18% in the exchange fraction calculated for individual aGUVs, a surprising result given previous studies using hemifusion reported considerably smaller uncertainties. These results highlight both the importance of interleaflet coupling in raft phenomena, and the urgent need to understand the sources of error that contribute to compositional uncertainty in aGUVs prepared by hemifusion.

Here, we extend our previous aGUV studies to cholesterol-containing bilayers that better mimic the plasma membrane composition of mammalian cells. We prepared aGUVs with inner leaflets composed of DPPC/16:1-PC/Chol 39/39/22 mol%, a mixture that exhibits Ld+Lo phase separation in symmetric model membranes at room temperature. The outer leaflets were hemifused to an SLB composed of 16:1-PC/Chol 80/20 mol%—a uniform, Ld mixture—and thus contain a greater proportion of unsaturated lipid compared to the inner leaflet. We observe both phase-separated and uniform aGUVs, with the latter having a greater degree of asymmetry on average, consistent with the presence of a phase boundary in the asymmetric composition space. We develop a coupled distributions model for determining the location of the phase boundary; the model accounts for uncertainty in the composition of individual aGUVs measured from changes in the fluorescence intensity of lipid probes that exchange between the GUV and SLB. We find that the magnitude of the compositional error is substantially greater than previously thought, and depends on whether the probe enters or exits the GUV during hemifusion.

## 2. Materials and Methods

### 2.1 Materials

Phospholipids 1,2-dipalmitoleoyl-sn-glycero-3-phosphocholine (16:1-PC) and 1,2-dipalmitoyl-sn-glycero-3-phosphocholine (DPPC) were purchased from Avanti Polar Lipids (Alabaster, AL) as a dry powder and used as supplied. Cholesterol (Chol) was purchased from Nu-Chek Prep (Elysian, MN) as a dry powder and used as supplied. Phospholipid and cholesterol stock solutions in HPLC-grade chloroform were prepared gravimetrically using an analytical balance with 0.1 mg precision. Fluorescent dyes were purchased from Avanti Polar Lipids (Alabaster, AL) as liquid stocks in chloroform and used as supplied: 1-palmitoyl-2-(dipyrrometheneboron difluoride)undecanoyl-sn-glycero-3-phosphocholine (TopFluor-PC or TFPC), 1,2-dioleoyl-sn-glycero-3-phosphoethanolamine-N-(lissamine rhodamine B sulfonyl) (ammonium salt) (Lissamine Rhodamine-PE or LRPE), and 1,2-dioleoyl-sn-glycero-3-phosphoethanolamine-N-(TopFluor AF594) (ammonium salt) (AF594). Dye stock solution concentrations were determined from absorbance measurements: TFPC was measured at 495 nm using an extinction coefficient of 96,904 M^-1^cm^-1^; LRPE was measured at 560 nm with an extinction coefficient of 95,000 M^-1^cm^-1^; and AF594 was measured at 590 nm with an extinction coefficient of 92,000 M^-1^cm^-1^. All stock solutions were stored at −20°C until use. Sucrose was purchased from Sigma-Aldrich and dissolved in ultrapure water to make a 100 mM stock solution. Sodium hydroxide was purchased from Fisher and used to make a 1 M cleaning solution. Sodium chloride was purchased from Sigma-Aldrich. 4-(2-hydroxyethyl)-1-piperazineethanesulfonic acid (HEPES), calcium chloride, and ethylenediaminetetraacetic acid (EDTA) were purchased from Thermo Scientific and used to make various buffer solutions.

### 2.2 Preparation of symmetric GUVs

Chloroform mixtures of lipids were prepared in glass culture tubes using a syringe and repeating dispenser (Hamilton USA, Reno, NV). Samples received a determined volume from chloroform lipid stocks to produce a composition of DPPC/16:1-PC/Chol = 39/39/22 mol% (250 nmol total lipid). The fluorescent probe TFPC, LRPE, or AF594 was added to achieve probe:lipid mol ratios of 1:1000, 1:5000, or 1:2500, respectively. The concentration of red dyes was kept low to minimize energy transfer from TFPC. GUVs were prepared by electroformation.^30^ Briefly, the chloroform solution of lipids and dye was deposited onto the ends of two ITO-covered slides. After evaporation of bulk chloroform, the slides were placed in a heated desiccator for 2 h at 55°C to remove residual solvent, resulting in a dry lipid film coating one end of the slides. A large O-ring was then pressed firmly onto the lipid film on one of the slides to create a chamber, and two small O-rings were placed on the clean end of the slide to act as spacers. After warm sucrose was added to the chamber, the second ITO slide was placed on top, and the assembly was stabilized with a clip. The slide “sandwich” was then placed into a pre-warmed metal block. A positive lead was attached to one slide and a negative lead to the other to create a capacitor. GUVs were produced by applying a 2 Vpp, 10 Hz sine wave for 2 h at 55°C. The block holding the sandwich was then allowed to cool to 23°C over 12 h. The GUVs were harvested and subsequently imaged with confocal fluorescence microscopy (CFM) as described below.

### 2.3 Preparation of asymmetric GUVs

#### 2.3.1 SLB formation

SLBs were prepared using previously described protocols with minor modifications.^23,31^ In brief, chloroform mixtures of lipids were prepared in glass culture tubes using a syringe and repeating dispenser. Samples received a determined volume from chloroform lipid stocks to produce a composition of 16:1-PC/Chol = 80/20 mol% (600 nmol total lipid). LRPE or TFPC was added to achieve a probe:lipid mol ratio of 1:10000 or 1:1000, respectively. This solution was used to prepare supported lipid bilayers (SLBs). 600 μL aqueous buffer (1 mM EDTA, 25 mM HEPES, 35 mM NaCl) was added to the chloroform lipid and dye mixture, and the suspension was subjected to vacuum while vortexing; this “rapid solvent exchange” (RSE) procedure removes chloroform and produces aqueous vesicles with a broad size distribution and low average lamellarity.^32–34^ The RSE suspension was then sonicated for 40 min in a cup horn sonicator (QSonica, Newtown, CT) at 50% power to produce small unilamellar vesicles (SUVs) of 75-100 nm diameter as verified by dynamic light scattering using a Litesizer 100 (Anton Parr, Graz, Austria). After sonication, the SUV solution was further diluted with 2.4 mL buffer to a final concentration of 0.2 mM. SLBs were prepared in 4-well chambered cover glass (ibidi, Gräfelfing, Germany) that was first cleaned with 1 M NaOH, copiously rinsed with ultrapure water, dried thoroughly at 55°C under vacuum for 3 min, and dusted with a nitrogen stream. The dish was then plasma cleaned (Harrick Plasma, Ithaca, NY) on the high setting for 2 min 45 s. Immediately after removing the dish from the plasma cleaner, the diluted SUV solution was deposited into the chambered cover glass with 750 μL pipetted into each well. The SLB dish was left at room temperature for 30 min, allowing time for the formation of an SLB as the SUVs adsorbed to the cover glass at the bottom of the chamber. The chamber was then flushed with ultrapure water to remove the remaining vesicles and imaged with CFM to check for quality, ensuring a flat SLB was present with no vesicles stuck to the surface.

#### 2.3.2 Hemifusion

aGUVs were prepared by calcium-induced hemifusion of the GUVs to the SLB, which joins their distal leaflets and allows for lipid exchange via diffusion^23^. GUVs were added to a well of the chambered cover glass that contained buffer (35 mM NaCl and 25 mM HEPES, pH 7.4) and the SLB. After allowing approximately 15 min for the GUVs to settle toward the SLB, a calcium-containing buffer (20 mM CaCl_2_, 25 mM HEPES, 5 mM NaCl, pH 7.4) was added for a final concentration of 5-6 mM CaCl_2_. After 15-30 min, an EDTA-containing buffer (30 mM EDTA, 25 mM HEPES, 5 mM NaCl, pH 7.4) was added to a final concentration of 6-7 mM EDTA in the well to chelate the calcium and terminate hemifusion. We caution that it is often necessary to adjust buffer concentrations to match osmolarities and thus prevent bursting of GUVs. Buffer osmolarities were checked using an OsmoTECH osmometer (Advanced Instruments, Norwood, MA) and adjusted as needed. After osmolarity matching, the EDTA-containing buffer typically had a diluted concentration of 20-25 mM EDTA. Lastly, the aGUVs were detached from the SLB by gentle pipet mixing with a large-orifice pipette tip.

One goal of our study was to assess the uncertainty in outer leaflet exchange fraction calculated from fluorescence intensity measurements. We performed the hemifusion experiment in two ways, demonstrated schematically in Fig. 1. In the “probe exit” experiment, GUVs containing TFPC were hemifused to an SLB to form aGUVs, and the amount of outer leaflet exchange was calculated from the decrease in TFPC fluorescence. In the “probe entry” experiment, GUVs were hemifused to a TFPC-containing SLB and the amount of exchange was calculated from the increase in TFPC fluorescence in the aGUV. In both cases, a trace amount of red fluorophore was included in either the SLB (for probe exit) or GUV (for probe entry) to aid in visualization. Because we used a very low concentration of the red probe to avoid fluorescence crosstalk, we did not calculate outer leaflet exchange fraction using the red signal. The calculations of exchange fraction using the TFPC signal are described in detail below.

**Figure 1.**
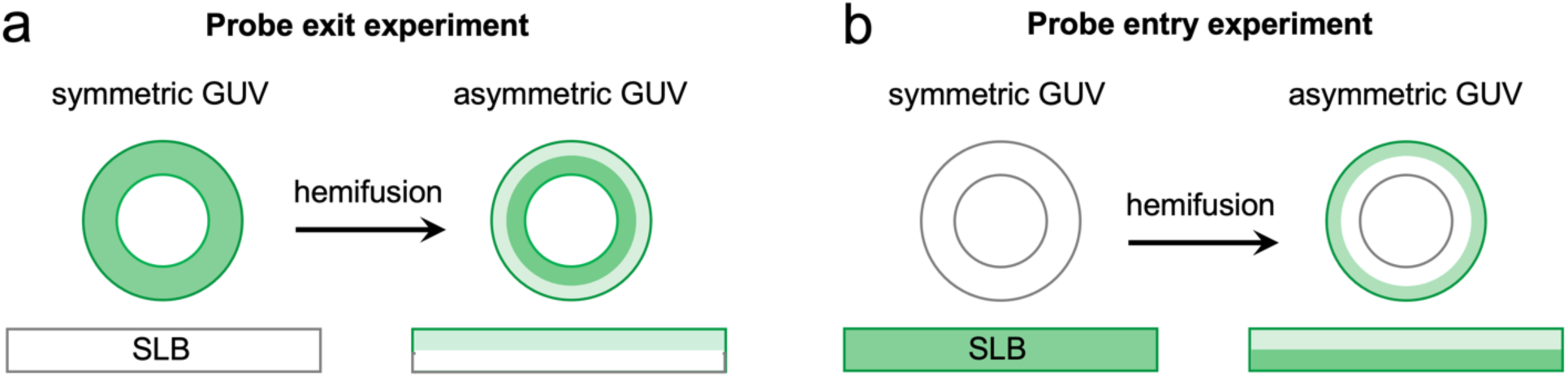
Schematic illustration of hemifusion experiments in this study. (a) In the probe exit experiment, the fluorescent lipid probe is initially present in the GUV. During hemifusion to an SLB that does not contain the probe, the intensity of the GUV decreases. (b) In the probe entry experiment, the fluorescent lipid is initially present only in the SLB. During hemifusion, intensity increases in the asymmetric GUV.

### 2.4 Imaging vesicles

All imaging was performed at 22°C, maintained with an objective cooling collar (Bioptechs, Butler, PA), using a Nikon C2+ point scanning system attached to a Nikon Eclipse Ti2-E microscope equipped with a Plan Apo Lambda 60x/1.4 NA oil immersion objective. We first imaged the symmetric GUVs in a well that contained an SLB and approximately 700 μL of buffer solution (25 mM HEPES, 35 mM NaCl, pH 7.4) to match the imaging conditions for aGUVs. Approximately 6 μL of the GUV solution was added to the well and allowed to settle for approximately 10 min. We looked for isolated, large, and round unilamellar vesicles to image. Once a suitable GUV was located, we centered it on the screen and set the zoom to 3x. The 488 nm laser was used to image GUVs containing TFPC while the 561 nm laser was used to image GUVs containing LRPE or AF594. For each image, the gain (HV) was set to 90 and the laser power was set to 7. Quarter waveplates (ThorLabs, Newton, NJ) were inserted in the excitation paths to correct for polarization artifacts as described previously^35^.

The procedure for imaging aGUVs was similar to that described above for symmetric GUVs with the following modifications. After the addition of EDTA-containing buffer and gentle pipette mixing to release aGUVs from the SLB, vesicles were allowed to settle for approximately 10 min. We used a macro to take sequential images in the red (561 nm laser, 546 nm quarter waveplate) and green (488 nm laser, 488 nm quarter waveplate) channels. The observation of fluorescence in both channels allowed us to determine whether a given GUV had exchanged lipid with the SLB.

### 2.5 Image analysis to determine the fraction of outer leaflet exchange

All images were saved as .nd2 files and subsequently processed in Fiji.^36^ To quantify lipid exchange, we analyzed only the green channel of GUVs imaged at the equator. The total fluorescence intensity was quantified in the same way for sGUVs prior to hemifusion and aGUVs after hemifusion. First, a circle was drawn on a GUV and the fluorescence intensity in the radial direction was extracted from the image in 1-degree increments (i.e., 360 values). The equatorial intensity *F* was calculated by averaging the maximum intensity values at each angle. The fraction of exchanged outer leaflet lipids, *ɛ_obs_*, for an individual aGUV was then calculated by comparing its total fluorescence *F*_*A*_ to the average initial fluorescence of sGUVs, *F̅_S_*. For probe exit experiments (i.e., where TFPC initially contained in sGUVs diffused into the SLB during hemifusion), *ɛ_obs_* is given by:

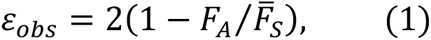

where *F̅_S_* is the average fluorescence intensity of *N_S_* symmetric vesicles prior to hemifusion:

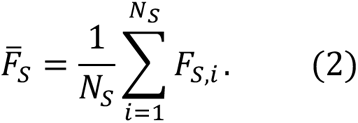

For probe entry experiments (i.e., where TFPC initially contained in the SLB diffused into GUVs during hemifusion), the fraction of outer leaflet exchange is given by:

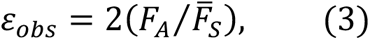

where *F̅*_*S*_ is the average fluorescence intensity of symmetric GUVs containing TFPC at a concentration identical to that of the SLB.

### 2.6 Exchange data analysis

#### 2.6.1 Coupled distributions model

Exchange data were analyzed with a coupled distributions model implemented in Mathematica v14.0 (Wolfram Research Inc., Champaign, IL) and described here. Hemifusion produces aGUVs of highly variable composition, ranging from no outer leaflet exchange to nearly complete exchange^23, 27–29^. To account for this variability, we use a modified exponential distribution to quantify the probability of obtaining an aGUV with true exchange fraction *ɛ* ∈ [0,1],

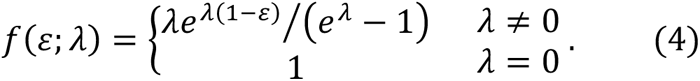

The parameter *λ* ∈ ℝ skews the distribution toward smaller (*λ* > 0) or larger (*λ* < 0) values of *ɛ*, with the special case *λ* = 0 corresponding to a uniform distribution in which all values of *ɛ* are equally likely.

The model must also account for the uncertainty in calculating *ɛ* from fluorescence intensity measurements that are subject to error. We introduce the notation *ɛ_obs_* to distinguish the observed exchange fraction of an aGUV (i.e., calculated from fluorescence measurements) from its true exchange fraction *ɛ*. Error propagation analysis of Eqs. 1 and 3 reveals that uncertainty in *ɛ_obs_* has a functional dependence on *ɛ* that is different for the probe exit and probe entry experiments (Supporting Section S1). Accordingly, we use normal distributions *g_exit_* and *g_entry_* to model the probability that an aGUV with true exchange value *ɛ* will produce a calculated exchange value *ɛ_obs_*, where the width of the distribution depends on *ɛ* according to Eq. S8a (for probe exit) or Eq. S8b (for probe entry):

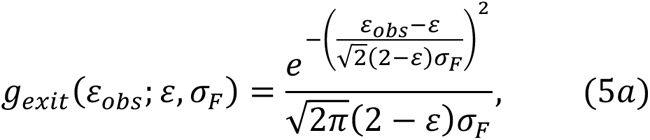

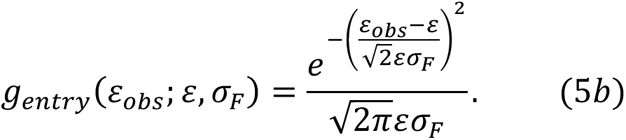

In Eqs. 5, *σ_F_* is a composite uncertainty that accounts for all sources of error inherent to measuring the average fluorescence intensity of a GUV from an equatorial confocal image.

The final addition to the model is the presence of a phase boundary, *ɛ*^∗^, in the asymmetric composition space, such that aGUVs with *ɛ* < *ɛ*^∗^ show phase coexistence, while aGUVs with *ɛ* > *ɛ*^∗^ are uniform. In cases where hemifusion produces a broad distribution of outer leaflet exchange that spans the phase boundary, two populations of vesicles (i.e., uniform and phase separated) will thus be observed. Combining Eqs. 4-5, the probability that a phase separated aGUV (i.e., for which 0 < *ɛ* < *ɛ*^∗^) will result in a measured exchange fraction 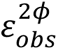 is

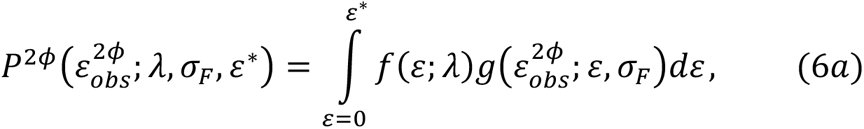

while the probability that a uniform aGUV (i.e., for which *ɛ*^∗^ < *ɛ* < 1) will result in a measured exchange fraction 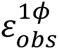 is

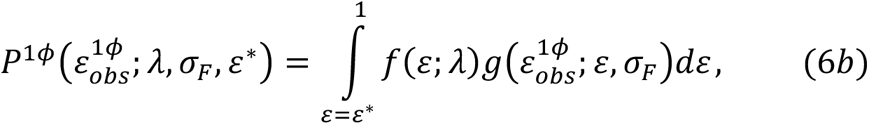

with *g* replaced by *g_exit_* (Eq. 5a) or *g_entry_* (Eq. 5b) for a probe exit or probe entry experiment, respectively. The probability distributions in Eqs. 6 are coupled by the shared parameters *ɛ*^∗^, *λ*, and *σ_F_*. The integrals do not have closed form solutions but can be numerically evaluated for data fitting.

#### 2.6.2 Data fitting

We denote the set of experimentally observed exchange fractions 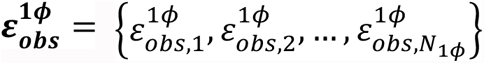 and 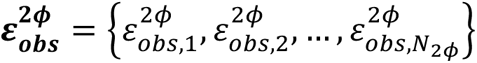 for uniform and phase separated aGUVs, respectively. The exchange datasets were binned and normalized to produce probability density histograms in which the total probability of all observations (i.e., uniform and phase separated) was unity. The normalized uniform and phase separated histograms were then jointly analyzed by fitting to Eqs. 6 using Mathematica’s NonlinearLeastSquares function with default settings. The adjustable parameters were *ɛ*^∗^, *λ*, and *σ_F_*.

## 3. Results

To model lipid asymmetry in the PM, we first prepared symmetric GUVs containing a mixture of low-melting lipid, high-melting lipid, and cholesterol, and therefore roughly mimicking the PM exofacial leaflet composition^37^. We then used hemifusion to effectively replace high-melting lipid from the GUV outer leaflet with low-melting lipid, consistent with the absence of high-melting lipids in the PM cytofacial leaflet^37^. In the resulting aGUVs, the leaflet identities are thus reversed compared to the PM (i.e., the aGUV inner leaflet mimics the PM exofacial leaflet, and the aGUV outer leaflet mimics the PM cytofacial leaflet). We chose DPPC and 16:1-PC as the high-melting and low-melting lipids, respectively. For the initial symmetric GUVs, we chose the composition DPPC/16:1-PC/Chol = 39/39/22 mol%, which exhibits liquid-disordered (Ld) + liquid-ordered (Lo) phase separation at 22°C. For the SLB, we used 16:1-PC/Chol = 80/20 mol%, a composition that is uniform at 22°C. We separately conducted probe-exit and probe-entry experiments using the same lipid probe, TFPC, which allowed us to evaluate differences in these methods of calculating exchange fraction.

### 3.1 Phase behavior of symmetric GUVs

The probe exit experiments consisted of 15 individual preparations and are summarized in Fig. S1 and Table S1. Electroformation at this composition resulted in a high yield of unilamellar vesicles with diameters greater than 10 μm, all of which were phase separated. The total intensity of individual vesicles (needed to calculate the fraction of outer leaflet exchange in aGUVs) is shown in Fig. S1A and was determined from equatorial confocal images as described in Methods. Within individual preparations, the relative standard deviation of the total intensity ranged from 9-26%, with an average relative standard deviation of 15% (Table S1).

Phase fractions of individual vesicles calculated from equatorial confocal images are shown in Fig. S1B. The average Ld area fraction varied from 0.42-0.55, with an overall mean of 0.52 and a standard deviation of 0.039 (Table S1). Assuming that the average equatorial phase fraction reflects that of the full GUV and using published values of Ld and Lo molecular areas for the similar system DPPC/DOPC/Chol (61.2 Å^9^/molecule and 43.5 Å^9^/molecule, respectively)^38^, we estimate the phase mole fractions to be *χ_Ld_* = 0.45 and *χ_Lo_* = 0.55.

The mean diameter of imaged GUVs across all preparations was 20 μm. GUVs as small as 4 μm and as large as 22 μm were included in subsequent analyses. We found little correlation between the vesicle size and total intensity (Fig. S2A) and a moderate positive correlation between Ld phase fraction and total intensity (Fig. S2B) that is consistent with the preference of TFPC for the Ld phase.

### 3.2 Phase behavior and composition of asymmetric GUVs measured from probe exit

We used calcium to induce hemifusion between symmetric DPPC/16:1-PC/Chol GUVs containing TFPC and an SLB composed of 16:1-PC/Chol 80/20 mol%, with a trace amount of LRPE included in the SLB to monitor exchange in real time. Fig. S3 shows several examples of the resulting aGUVs imaged in both the red and green channels, with the presence of LRPE confirming the successful exchange of lipid between the SLB and GUV. The fraction of exchanged lipid, *ɛ_obs_*, was calculated from the decrease in TFPC intensity as described in Methods. Consistent with previous reports, *ɛ_obs_* varied substantially between aGUVs as shown in Fig. 2a and covered the full range of values expected for hemifusion (i.e., 0 < *ɛ_obs_* < 100%). Values lying outside the expected range were also obtained, most likely due to errors inherent in the calculation of *ɛ*_*obs*_ discussed in detail below. Such values can also occur when vesicles have multiple lamellae (*ɛ_obs_* < 0%) or undergo full fusion rather than hemifusion (*ɛ_obs_* > 100%). Representative images of aGUVS with different amounts of outer leaflet exchange are shown in Fig. 3.

**Figure 2.**
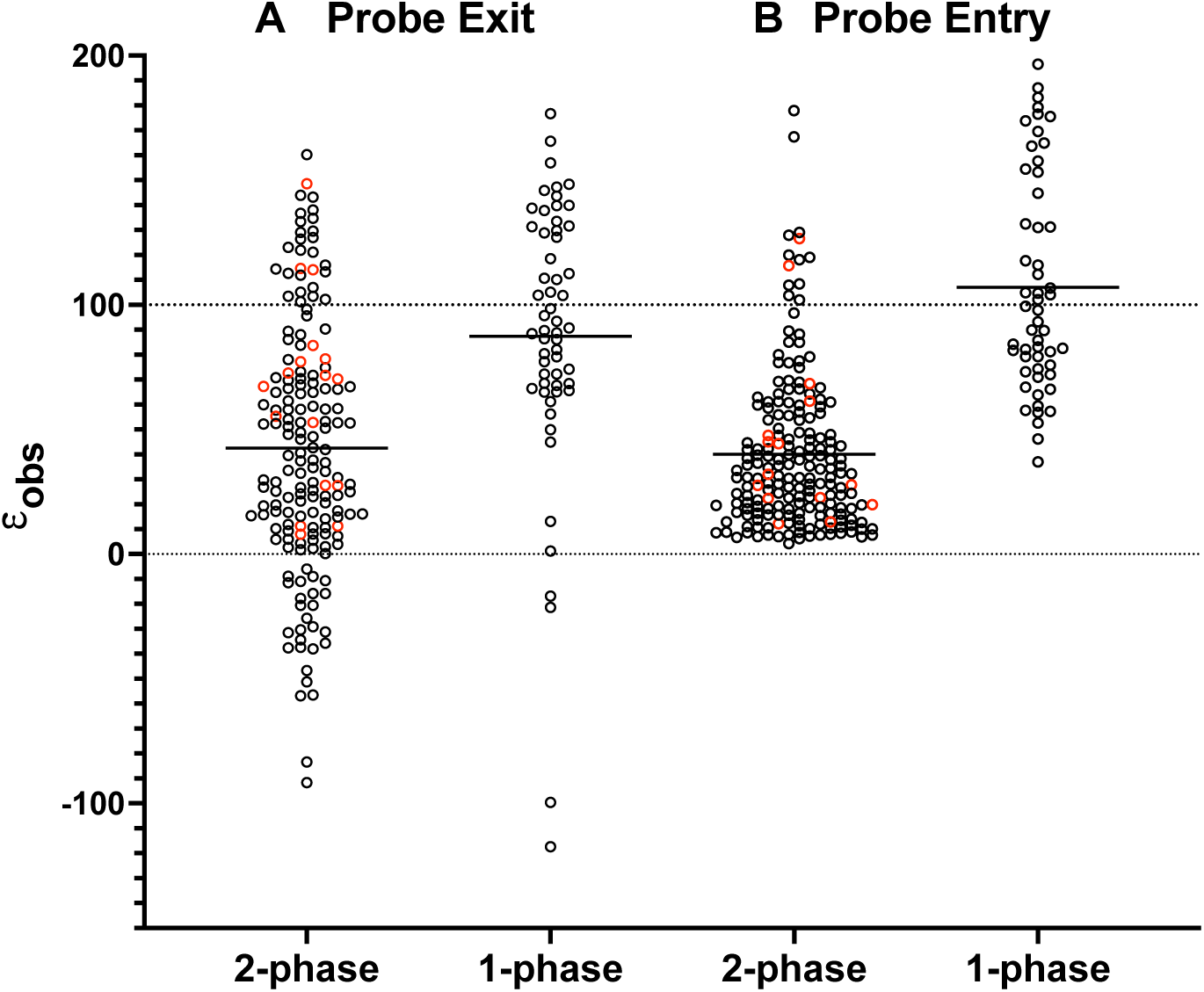
Exchange fraction of aGUVs calculated from probe exit and entry experiments. Distributions of exchange percentage *ɛ*_*obs*_ calculated from phase separated and uniform aGUVs in the probe exit (left two columns) and probe entry (right two columns) experiments. Red circles represent aGUVs showing modulated phase patterns and solid horizontal lines represent the mean of the distribution. Dotted lines at 0 and 100% denote the limiting values when only one leaflet undergoes exchange, as expected for hemifusion.

**Figure 3.**
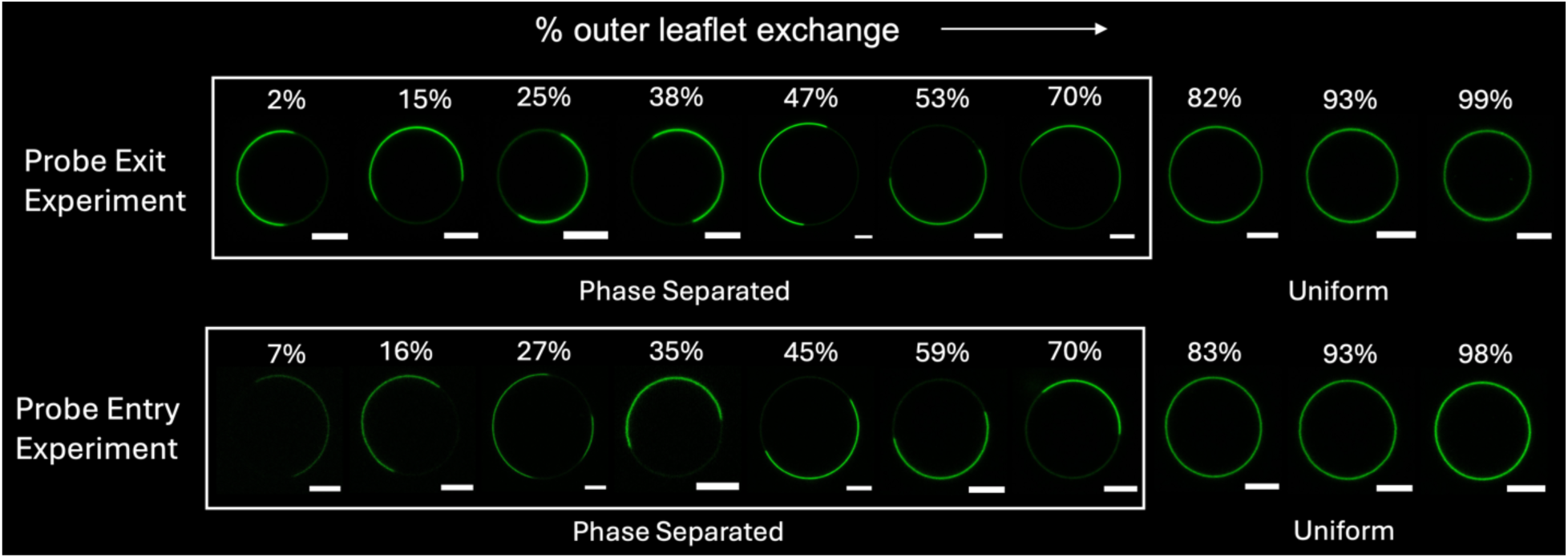
Representative aGUVs at different levels of exchange. aGUVs obtained from the probe exit experiment (top row) and probe entry experiment (bottom row) with different calculated exchange percentage as indicated. Scale bar 5 μm.

After hemifusion we observed a substantial population of uniform (i.e., one-phase) aGUVs in addition to a larger population of aGUVs that, like the initial symmetric GUVs, were phase separated (Fig. 2 and Fig. 3). Comparing the phase separated aGUVs from the different preparations, the average Ld area fraction was similar to that of the initial symmetric GUVs, ranging from 0.45-0.55, with an overall mean of 0.51 and a standard deviation of 0.026 (Fig. S4 and Table S2). The average outer leaflet exchange fraction of the phase separated aGUVs (*ɛ̅_obs_* = 0.43, *σ* = 0.50, *N* = 172) was about half that of the uniform aGUVs (*ɛ̅_obs_* = 0.87, *σ* = 0.57, *N* = 57). The difference between these mean values, 0.45 ± 0.17 (95% confidence interval), is statistically significant (*p* < 0.0001). These observations are consistent with the presence of a phase boundary in the asymmetric composition space parameterized by *ɛ*; here and throughout, we refer to the location of this boundary as *ɛ*^∗^. We conclude that asymmetric GUVs with a true exchange fraction *ɛ* < *ɛ*^∗^ are phase separated in at least one leaflet, while those with *ɛ* > *ɛ*^∗^ have crossed a phase boundary into a region of the asymmetric composition space where both leaflets are uniformly mixed. Notably, the distributions of phase separated and uniform vesicles in Fig. 2a show substantial overlap, consistent with a large uncertainty in calculated exchange fraction, which we discuss below.

### 3.3 Phase behavior and composition of asymmetric GUVs measured from probe entry

In previous work, we found that exchange fractions calculated from entry of TFPC showed smaller errors compared to those calculated from exit of DiD^28^. It is unclear if this difference is mainly related to the mode of exchange, or to the different probes that were used. To address this question, we conducted 12 probe entry experiments in which symmetric GUVs that lacked TFPC were hemifused to an SLB that contained TFPC. *ɛ_obs_* was calculated from the increase in TPFC intensity in the aGUV as described in Methods. The combined results of these experiments are shown in Fig. 2b. As was the case for probe exit experiments (Fig. 2a), we observed both uniform and phase separated aGUVs. Comparing the different preparations, the average Ld area fraction of phase separated aGUVs ranged from 0.47-0.62, with an overall mean of 0.54 and a standard deviation of 0.045 (Fig. S5 and Table S3). The average outer leaflet exchange fraction calculated for uniform aGUVs (*ɛ̅_obs_* = 1.07 *σ* = 0.43, *N* = 54) was again significantly larger (*p* < 0.0001) than that calculated for phase separated aGUVs (*ɛ̅_obs_* = 0.40, *σ* = 0.31, *N* = 193). We note that, to calculate the exchange fraction via Eq. 3, it is necessary to measure TFPC intensity in symmetric GUVs at its nominal concentration in the SLB; these data are shown in Fig. S6 and summarized in Table S4.

### 3.4 Location and uncertainty of the asymmetric phase boundary

The substantial overlap in the distributions of uniform and phase separated aGUVs shown in Fig. 2 makes it difficult to determine the location of the phase boundary (*ɛ*^∗^) by simple visual inspection. To objectively determine both *ɛ*^∗^ and its uncertainty, we developed a coupled distributions model to fit the experimental exchange data, described in Methods and Supporting Section S1. The salient features of the model are demonstrated in Fig. 4a-d and summarized here:

1. The probability *f*(*ɛ*) for an aGUV to have a true exchange fraction *ɛ* is assumed to be a function of a single adjustable parameter, *λ*, according to Eq. 4. When *λ* = 0, all values of *ɛ* are equally likely. Positive (negative) values of *λ* shift the distribution toward smaller (larger) values of *ɛ* (Fig. 4a).
2. The true exchange fraction distribution *f*(*ɛ*) is smeared by the uncertainty associated with calculating the exchange fraction from confocal fluorescence images via Eqs. 1 and 3. We account for this smearing with a normal distribution, *g*(*ɛ_obs_*), that describes the probability of calculating an exchange fraction *ɛ_obs_* for an aGUV in a probe exit (Eq. 5a) or probe entry (Eq. 5b) experiment, given that the aGUV has a true exchange fraction *ɛ*. Error propagation analysis of Eqs. 1 and 3 reveals that the widths of these distributions, 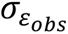 depend on both *ɛ* and the relative error associated with measuring aGUV fluorescence intensity, *σ_F_*, as shown in Fig. 4b. In the probe exit experiment, 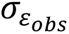 is largest at low levels of exchange and decreases linearly with increasing *ɛ*. In the probe entry experiment, this trend is reversed: error is smallest at low levels of exchange and increases linearly with increasing *ɛ*.
3. When the asymmetric composition space contains a phase boundary *ɛ*^∗^, the probability distributions of *ɛ_obs_* for uniform and phase separated vesicles (*P*^1*ϕ*^ and *P*^2*ϕ*^, respectively) are given by Eqs. 6. These distributions are uniquely determined by the shared parameters *λ*, *σ_F_*, and *ɛ*^∗^, and are thus intrinsically coupled. Fig. 4c-d demonstrates how *P*^1*ϕ*^ and *P*^2*ϕ*^ spread out with increasing *σ_F_* for the probe exit (Fig. 4c) and probe entry (Fig. 4d) experiments.

**Figure 4.**
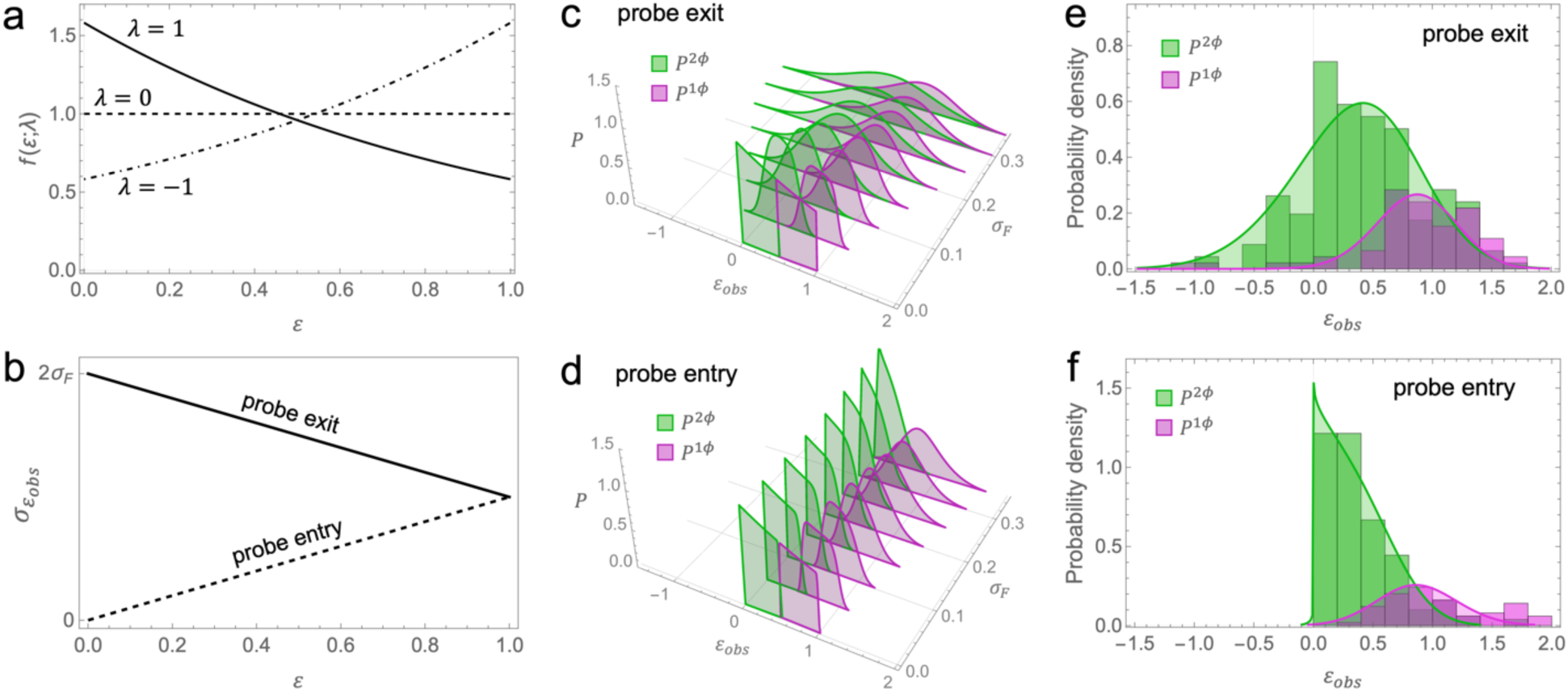
Coupled distributions model. (a) Model for the probability distribution of true exchange fraction *f*(*ɛ*) (Eq. 4) shown for three values of the parameter *λ*. (b) Theoretical dependence of uncertainty in calculated exchange fraction 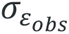 vs. true exchange fraction *ɛ* for probe exit (solid line) and probe entry (dashed line) experiments derived from error propagation analysis. (c and d) Model for the coupled probability distributions *P*^2*ϕ*^(*ɛ_obs_*) and *P*^1*ϕ*^(*ɛ_obs_*) (green and pink, Eqs. 6a and 6b, respectively) for the probe exit (c) and probe entry (d) experiments, calculated for different values of the parameter *σ_F_* and fixed values *ɛ*^∗^ = 0.5 and *λ* = 1. (e and f) Histograms of probe exit (e) and probe entry (f) data overlaid with the best fit curves from fitting to the coupled distributions model. Best fit parameters are given in Table 1.

**Table 1.** Best fit parameters from the coupled distributions model for probe entry and exit.

| Parameter | Probe exit | Probe entry | Joint analysis |
| --- | --- | --- | --- |
| $N_{1\phi}$ | 172 | 193 | |
| $N_{2\phi}$ | 57 | 54 | |
| $\varepsilon^*$ | $0.72 \pm 0.05$ | $0.76 \pm 0.03$ | $0.75 \pm 0.02$ |
| $\sigma_F$ | $0.29 \pm 0.02$ | $0.38 \pm 0.06$ | $0.31 \pm 0.02$ |
| $\lambda$ | $0.5 \pm 0.5$ | $0.3 \pm 0.3$ | $0.4 \pm 0.3$ |

The exchange data from Fig. 3 were analyzed by binning the *ɛ_obs_* values for uniform and phase separated aGUVs and then fitting the resulting histograms to the coupled distributions model. Separate fits were performed for the probe exit and probe entry data, with the results shown in Fig. 4e (probe exit) and Fig. 4f (probe entry) and summarized in Table 1. The phase boundaries determined from the probe exit and probe entry data (*ɛ*^∗^ = 0. 72 ± 0.05 and 0. 76 ± 0.03, respectively) agree to within experimental uncertainty. The best fit values of *λ* (0. 5 ± 0.5 and 0. 3 ± 0.3 for probe exit and entry, respectively) are also similar and indicate that outer leaflet exchange is essentially uniformly distributed for our experimental conditions, possibly with a slight preference for lower amounts of exchange. Interestingly, the values of *σ_F_* obtained from model fitting (0. 29 ± 0.02 and 0. 38 ± 0.06 for probe exit and entry, respectively) were substantially larger than implied by fluorescence measurements of symmetric GUVs. In the Discussion, we speculate on potential reasons for this discrepancy.

Given the overall similarity of fit parameters obtained from the separate analyses of probe exit and entry data, we also performed a joint refinement to a single global parameter set, which yielded best fit values of *ɛ*^∗^ = 0. 75 ± 0.02, *λ* = 0.4 ± 0.3, and *σ_F_* = 0.31 ± 0.02 (Table 1).

## 4. Discussion

### 4.1 Interleaflet coupling can disrupt liquid-liquid phase separation in asymmetric GUVs

Our experimental system consisted of an initially phase separated composition, DPPC/16:1-PC/Chol = 39/39/22 mol% at 22°C, that is far from a phase boundary in composition and temperature. Symmetric GUVs prepared at this composition were phase separated at 22°C, with approximately equal area fractions of Ld and Lo phase. After exchange of outer leaflet lipids by calcium-induced hemifusion, a substantial fraction of the resulting asymmetric GUVs showed uniform fluorescence, indicating a single phase. The average exchange fraction of the uniform aGUVs was significantly larger than that of phase separated aGUVs, consistent with the presence of a phase boundary in the asymmetric composition space of this mixture. Stated another way, Lo domains in the inner leaflet were disrupted by exchanging DPPC for 16:1-PC in the outer leaflet. We estimate the outer leaflet composition at the phase boundary to be DPPC/16:1-PC/Chol = 9/70/21 mol%. In fully symmetric bilayers, this composition is in close proximity to an Ld + Lo phase boundary (Fig. S7), suggesting that domain disruption in the aGUV inner leaflet closely coincides with the loss of a driving force for lipid segregation in the outer leaflet.

Our finding of a phase boundary in the asymmetric composition space is a striking example of a type of interleaflet coupling that London has termed “uniform leaflet dominance”^39^, and that has previously been observed in asymmetric planar supported bilayers^40^ and unsupported bilayers^41^, as well as large unilamellar vesicles prepared by cyclodextrin-mediated exchange^39^. We previously reported uniform leaflet dominance in DPPC/DOPC aGUVs exhibiting gel + fluid phase separation^28^. To our knowledge, the present study is the first to show uniform leaflet dominance in aGUVs with Ld + Lo phase separation. Uniform leaflet dominance occurs when the unfavorable mismatch free energy of anti-registered domains (i.e., Ld directly opposite Lo) is so large that it overwhelms the favorable change in free energy contributed by lateral phase separation^9^. Simple theoretical treatments can relate the location of the phase boundary *ɛ*^∗^ to the strength of in-plane vs. midplane lipid interactions^42^. Planned studies using different low-melting and high-melting lipids may shed light on the molecular determinants of interleaflet coupling.

### 4.2 The highly variable asymmetry of hemifusion vesicles is useful

The use of calcium-induced hemifusion to prepare asymmetric giant vesicles results in wide vesicle-to-vesicle variability in lipid asymmetry within a population of aGUVs, as shown in Figs. 2 and 3. Enoki and Feigenson have noted the advantages of quantifying asymmetry of individual GUVs, in particular emphasizing the ability to exclude vesicles that are not sufficiently asymmetric from subsequent analysis^23^. The choice to focus on highly asymmetric GUVs is understandable from a biological standpoint, given the strong asymmetries (e.g., in lipid headgroups and chain unsaturation) known to exist in the PM^13^.

While we agree that quantifying the asymmetry of individual aGUVs is essential, and that highly asymmetric vesicles hold special significance, we argue that the large variability in outer leaflet exchange inherent to the hemifusion method is more feature than bug. First, analyzing all aGUVs—including those with lesser amounts of exchange—provides an opportunity to elucidate the dependence of any measurable membrane property (e.g., phase state, order, permeability, bending stiffness, etc.) on the extent of asymmetry, and thus potentially shed light on the nature of interleaflet coupling. Second, it is unlikely that the highly asymmetric PM composition reported in Lorent et al.^13^ is the only physiologically relevant state. Indeed, cells can release PM asymmetry through the activation of scramblases^43, 44^, which suggests that the degree of membrane asymmetry might be a continuous variable under dynamic cellular control. In this light, the individual aGUVs in a hemifusion experiment, when sorted by the amount of outer leaflet exchange, represent a series of snapshots along a path of increasing asymmetry. Together, they provide a window into how membrane properties change as the lipids in one leaflet are gradually replaced by a different set of lipids.

### 4.3 Compositional uncertainty of aGUVs is different for probe entry vs. exit

This work improves upon previous efforts to estimate the compositional uncertainty of aGUVs prepared by hemifusion. We assume that the error is dominated by vesicle-to-vesicle variation in fluorescence intensity that occurs even in nominally identical GUVs, and that arises from sources including instrumentation (e.g., spatial and temporal fluctuations in illumination intensity)^45^, probe photostability, and compositional variability^46^. Phase separated vesicles are further subject to differences in the apparent phase fraction seen in different confocal slices. Since aGUVs are also prone to each of these sources of uncertainty in addition to others discussed below, vesicle-to-vesicle variation sets a lower limit for uncertainty when calculating exchange from Eqs. 1 and 3. The data in Tables S1 and S4 thus provide lower-bound estimates for *σ_F_* of 10-25%.

A recent study from our group used an empirical maximum likelihood approach to estimate the compositional uncertainty of binary DPPC/DOPC aGUVs^28^. In that case, symmetric GUVs that initially showed gel + fluid phase separation became homogeneously mixed when approximately 60% of the outer leaflet DPPC was replaced with DOPC, resulting in two vesicle populations (i.e., phase separated and uniform). Similar to the present study, some overlap in the distributions was observed, but the number of data points was too small to produce robust histograms for data fitting. Instead, the number of vesicles in the overlapping region was compared to simulated distributions to determine the most likely value of 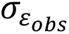, which was found to be 18% for the exiting probe and 12% for the entering probe. This analysis did not attempt to account for how the uncertainty might depend on the mode of probe exchange (i.e., exit vs. entry) or extent of exchange. It was also assumed that all values of exchange are equally probable, which is unlikely to generally be true, though experimental conditions can clearly be found such that this assumption is approximately met.

Here, we developed a quantitative model to address the shortcomings mentioned above. Our starting point was an error propagation analysis of the equations used to calculate exchange fraction from GUV intensities. This analysis led to several surprising predictions, summarized graphically in Fig. 4b: (1) when exchange fraction is calculated from the decrease in intensity of a probe that exits the GUV during hemifusion, the associated error will *decrease* with increasing amount of exchange; (2) the opposite trend will be observed when exchange is calculated from a probe that enters the GUV from the SLB during hemifusion (i.e., error will *increase* with increasing amounts of exchange); (3) if the uncertainty associated with a GUV fluorescence measurement, *σ_F_*, is independent of the mode of probe transfer (a reasonable assumption), then the entry experiment will always yield more precise estimates of exchange than the exit experiment. The coupled distributions model incorporates these predictions of error propagation and thus provides a means of testing them. Indeed, we find that the experimental distributions are well described by the model using the parameters in Table 1, lending support to our hypotheses about the underlying compositional uncertainty. It is also notable that Enoki and Heberle^28^ reported a value of 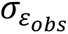 for the exiting probe that is 1.5-fold greater than that of the entering probe, in qualitative agreement with our prediction.

We were surprised by the magnitude of *σ_F_* obtained from fitting the distributions (30-40%, Table 1), which was substantially greater than the lower-bound estimate based on the spread in fluorescence intensity seen in symmetric vesicles (10-25%, Tables S1 and S4). As mentioned above, a major difference between symmetric and asymmetric vesicles is that aGUVs prepared by hemifusion have a greater intrinsic compositional variation than conventionally prepared symmetric GUVs. Because probe intensity is often strongly dependent on composition due to fluorophore environmental sensitivity^47, 48^, the large compositional variability of aGUVs likely contributes to a greater variability in probe intensity. Another confounding factor is the possibility of full fusion, which was previously observed in some vesicles using similar calcium concentrations^23^. Values of *ɛ_obs_* greater than 1 and as high as 2 would be expected when both GUV leaflets are available for exchange with the SLB. The presence of MLVs would have the opposite effect, resulting in *ɛ_obs_* less than zero and as low as −1 in probe exit measurements.

The values for 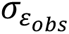 obtained from our analysis are substantially larger than values derived from maximum likelihood estimation as discussed previously^28^. An additional factor that contributes to greater uncertainty in the present study is our use of a single probe to calculate *ɛ_obs_*. In previous studies, different color fluorophores were included in the pre-hemifusion GUVs and SLB, thus enabling simultaneous and independent measurements of *ɛ_obs_* from probe exit and entry^23, 27–29^. Typically, the average of two independent measurements of comparable error will reduce the uncertainty by a factor of √2. However, as described above, propagation of uncertainty reveals that the error associated with probe entry is generally smaller than that of probe exit at all values of exchange (as in Fig. 4b), assuming that *σ_F_* is independent of the mode of probe transfer. In this case, the uncertainty associated with the average of the two measurements will be greater than that of the probe entry measurement alone until the exchange fraction reaches a value of 0.73. It is therefore unlikely that smaller uncertainties reported in previous hemifusion studies can be explained by averaging of the probe exit and probe entry values. Instead, we believe the smaller errors are related to the use of selection criteria: specifically, aGUVs were excluded from subsequent analysis if the value of *ɛ_obs_* measured from exiting and entering probes disagreed by more than a specified threshold (e.g., 20%), or if either *ɛ_obs_* value was deemed unphysical (i.e., less than 0 or greater than 100% exchange)^23, 27–29^. We speculate that applying these criteria may have the effect of eliminating MLVs or fully fused vesicles that are otherwise difficult to identify. We chose to include all vesicles in our analysis with the goal of determining the true uncertainty of exchange measurements. This information, and the methodology for determining error introduced here, should prove valuable in the continued development and optimization of the hemifusion technique.

## Author contributions

F.A.H. conceived of the project and obtained funding. K.B.K. and F.A.H. designed the research. K.B.K. performed the experiments. K.B.K. and F.A.H. analyzed the data and wrote the manuscript.

## Acknowledgements

We thank Dr. Thais Enoki and Dr. Haden Scott for help with the hemifusion protocol. This research was supported by NIH grant R01 GM138887 (to F.A.H). K.B.K. was supported by the Graduate Advancement & Training Education (GATE) fellowship from the University of Tennessee-Oak Ridge Innovation Institute (UT-ORII).

