## Supporting information for "Disruption of liquid/liquid phase separation in asymmetric GUVs prepared by hemifusion"

*corresponding author

**This PDF file includes:**

Figures S1-S7
Tables S1-S4
Section S1

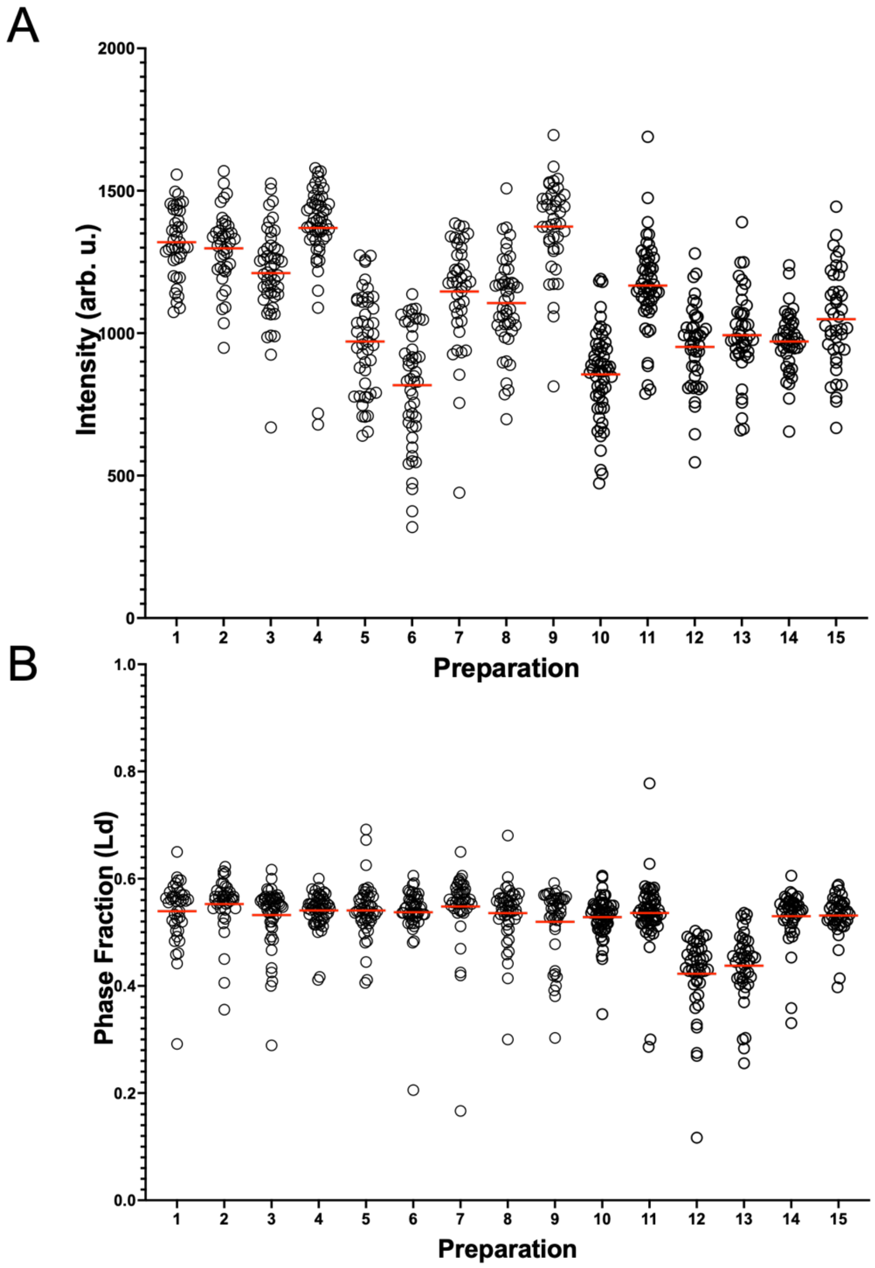


**Figure S1.** Distributions of vesicle intensity (A) and Ld phase fraction (B) of symmetric (i.e., pre-hemifusion) GUVs for the probe exit experiment, separated by individual preparation. All vesicles were phase separated. Horizontal lines represent the mean intensity for each preparation.

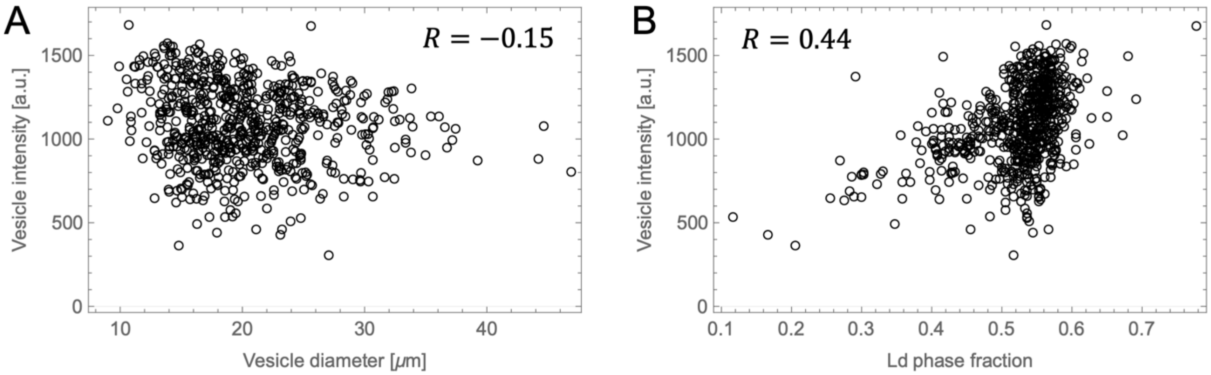


**Figure S2.** A, vesicle intensity and vesicle diameter show a weak negative correlation (Pearson correlation coefficient $R = -0.15$). B, vesicle intensity and Ld phase fraction show a moderate positive correlation ($R = 0.44$).

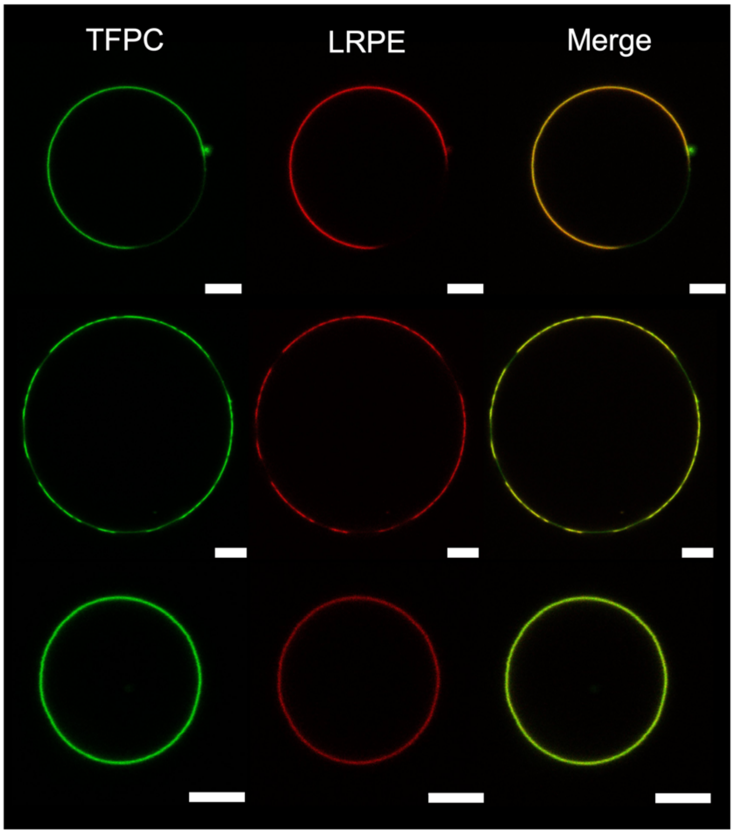


**Figure S3.** Examples of DPPC/16:1-PC/Cholesterol aGUVs prepared by calcium-induced hemifusion, and using the probe exit mode of exchange. The initial symmetric GUVs contained only TFPC. The aGUVs also contain LRPE that was incorporated into the SLB. Scale bar is 5 µm.

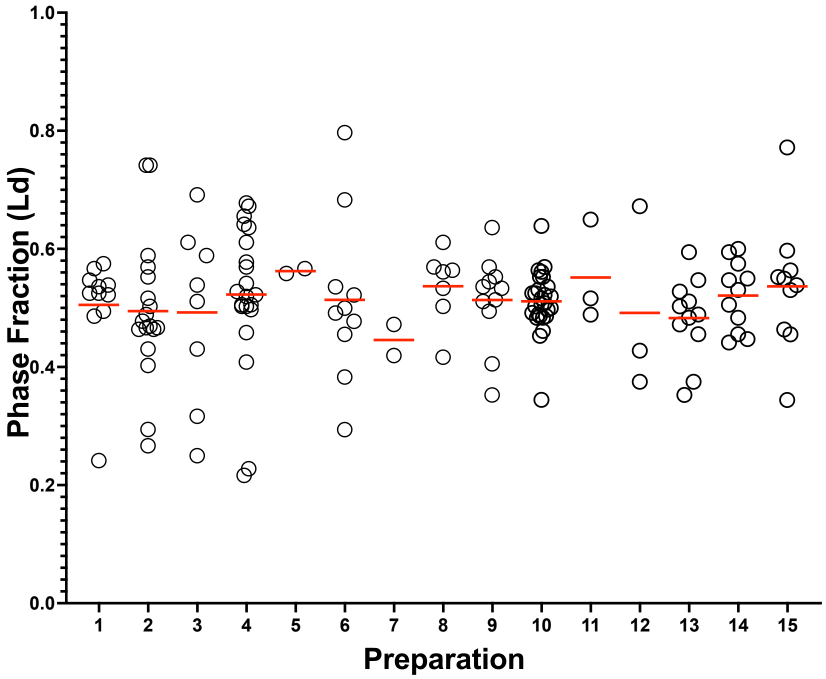


**Figure S4.** Ld phase fractions for each phase separated aGUV from the probe exit experiment separated by individual preparation. Horizontal lines represent the mean Ld phase fraction for each preparation.

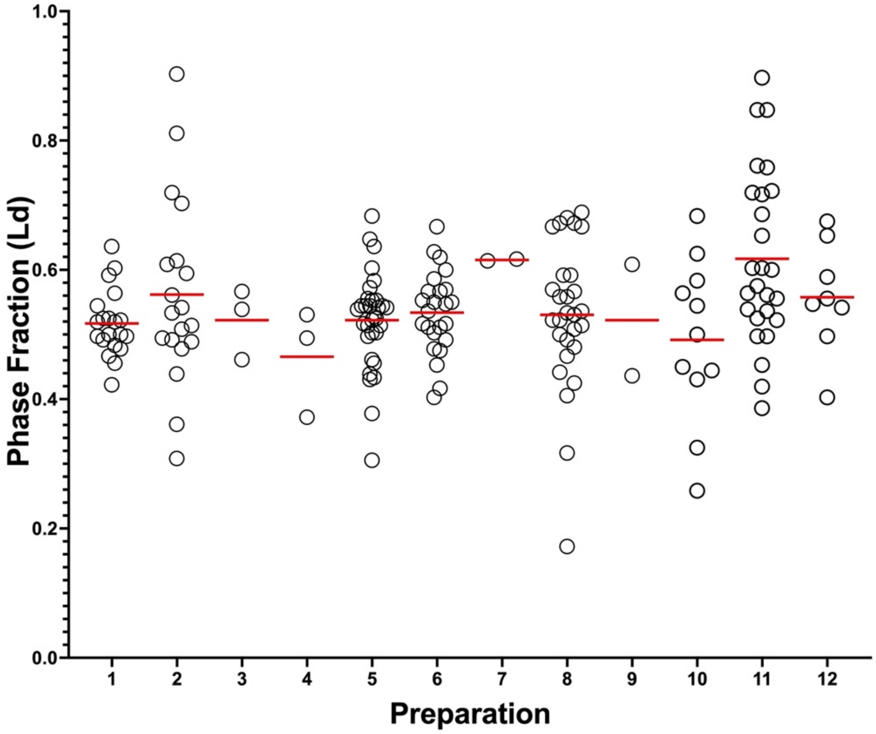


**Figure S5.** Ld phase fractions for each phase separated aGUV from the probe entry experiment separated by individual preparation. Horizontal lines represent the mean Ld phase fraction for each preparation.

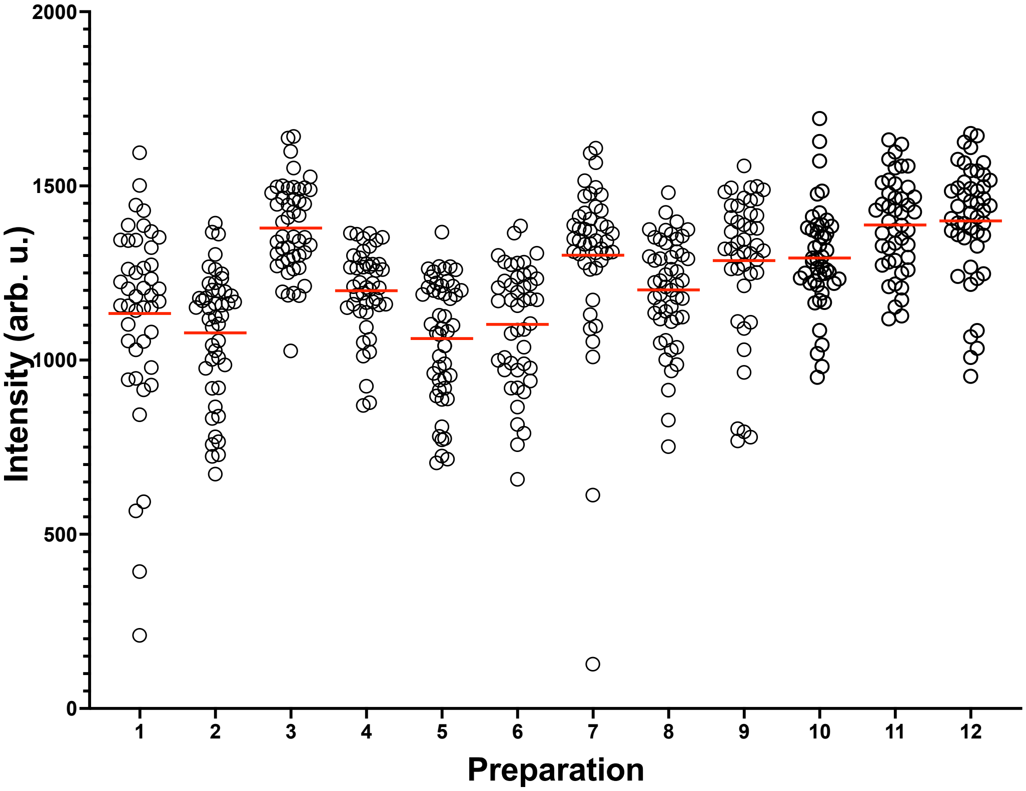


**Figure S6.** Control sGUVs for the probe entry experiment separated by individual preparation plotted with their respective intensities. All vesicles were phase separated. Horizontal lines represent the mean intensity for that preparation.

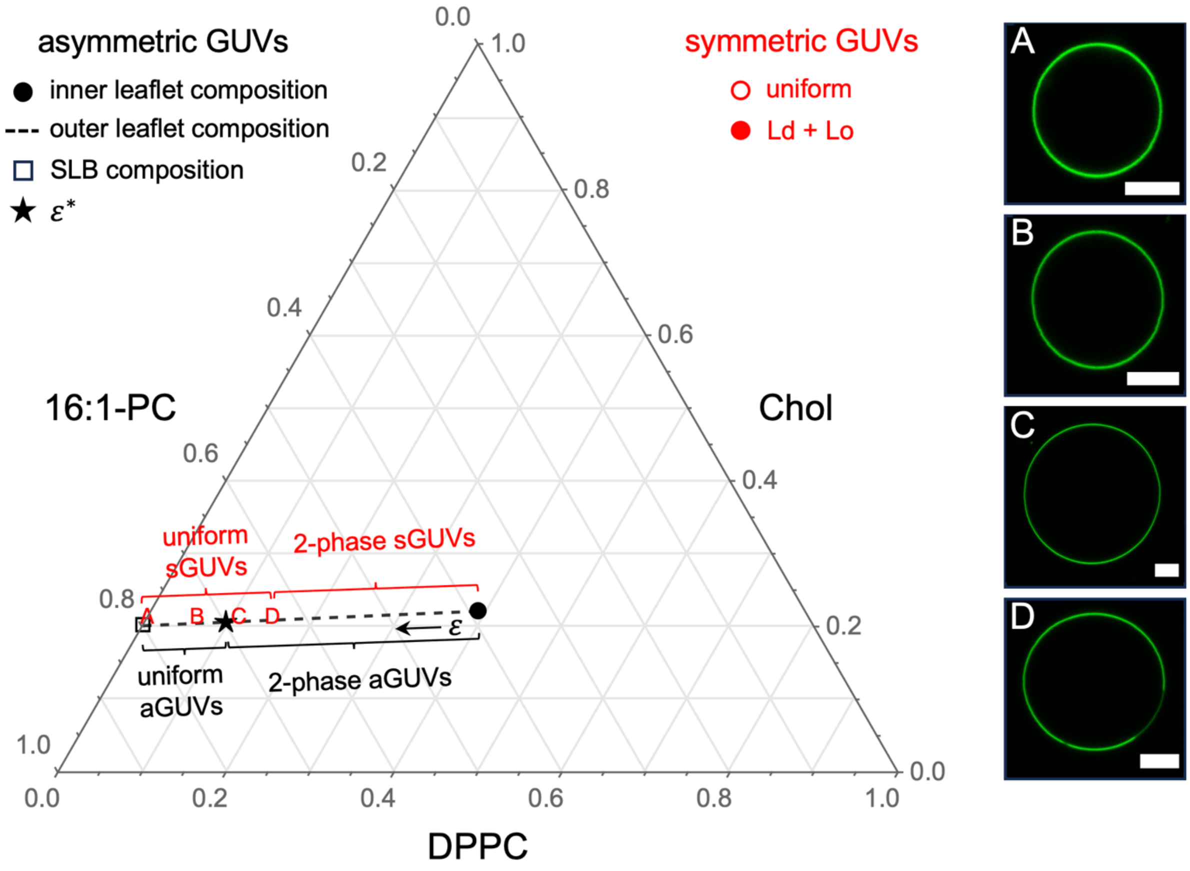


**Figure S7 Comparing the phase behavior of symmetric and asymmetric GUVs.** The aGUV inner leaflet composition is marked by the solid black circle. aGUV outer leaflet compositions fall along the dashed line joining the inner leaflet composition with the SLB composition (open square). The direction of increasing outer leaflet exchange, $\varepsilon$ , is shown with an arrow, and the asymmetric phase boundary ($\varepsilon^* = 0.75$) is marked by a solid black star. Shown on the right side are images of symmetric GUVs A-D with compositions marked by red symbols on the phase diagram. Compositions A-C produced uniform GUVs while composition D showed coexisting Ld + Lo phases.

**Table S1.** Data for symmetric (pre-hemifusion) GUVs used in probe exit experiments: number of vesicles ($N$), average vesicle intensity ($\bar{F}$), relative uncertainty in intensity ($\sigma_{\bar{F}}/\bar{F}$), average diameter ($d$), Ld phase area fraction ($f_{Ld}$), relative uncertainty in Ld area fraction ($\sigma_{f_{Ld}}/f_{Ld}$).

| Prep | $N$ | $\bar{F}$ | $\sigma_{\bar{F}}/\bar{F}$ (%) | $d$ ($\mu m$) | $f_{Ld}$ | $\sigma_{f_{Ld}}/f_{Ld}$ (%) |
|---|---|---|---|---|---|---|
| DOPC | 30 | 1040 ± 106 | 10.2 | 20 ± 6 | -- | -- |
| 1 | 40 | 1320 ± 122 | 9.2 | 19 ± 4.4 | 0.54 ± 0.059 | 11.0 |
| 2 | 40 | 1300 ± 130 | 10.0 | 27 ± 6.1 | 0.55 ± 0.051 | 9.2 |
| 3 | 50 | 1210 ± 155 | 12.8 | 23 ± 4.5 | 0.53 ± 0.057 | 10.8 |
| 4 | 53 | 1370 ± 168 | 12.3 | 19 ± 4.2 | 0.54 ± 0.033 | 6.2 |
| 5 | 47 | 971 ± 174 | 18.0 | 21 ± 4.8 | 0.54 ± 0.051 | 9.4 |
| 6 | 47 | 817 ± 209 | 25.6 | 19 ± 4.8 | 0.54 ± 0.056 | 10.4 |
| 7 | 42 | 1150 ± 188 | 16.4 | 20 ± 5 | 0.55 ± 0.073 | 13.4 |
| 8 | 47 | 1110 ± 165 | 14.8 | 18 ± 5.1 | 0.54 ± 0.056 | 10.4 |
| 9 | 42 | 1370 ± 161 | 11.7 | 16 ± 3.6 | 0.52 ± 0.067 | 12.9 |
| 10 | 57 | 852 ± 158 | 18.5 | 19 ± 4.5 | 0.53 ± 0.039 | 7.4 |
| 11 | 53 | 1170 ± 159 | 13.6 | 26 ± 3.5 | 0.54 ± 0.065 | 12.1 |
| 12 | 43 | 952 ± 146 | 15.3 | 20 ± 4 | 0.42 ± 0.074 | 17.4 |
| 13 | 42 | 993 ± 154 | 15.5 | 18 ± 4.0 | 0.44 ± 0.064 | 14.7 |
| 14 | 42 | 971 ± 108 | 11.1 | 19 ± 4.0 | 0.53 ± 0.050 | 9.4 |
| 15 | 41 | 1050 ± 174 | 16.5 | 19 ± 5.0 | 0.53 ± 0.038 | 7.19 |
| Mean | 46 | 1110 ± 182 | 14.8 ± 4.1 | 20 ± 3 | 0.52 ± 0.039 | 10.8 ± 3.0 |

**Table S2.** Data for aGUVs obtained from probe exit experiments: number of vesicles ($N$), number of phase separated vesicles including modulated ($N_{2\phi}$), number of modulated vesicles ($N_{mod}$), Ld phase area fraction ($f_{Ld}$), relative uncertainty in Ld area fraction ($\sigma_{f_{Ld}}/f_{Ld}$).

| Prep | $N$ | $N_{2\phi}$ | $N_{mod}$ | $f_{Ld}$ | $\sigma_{f_{Ld}}/f_{Ld}$ (%) |
|---|---|---|---|---|---|
| 1 | 20 | 11 | 0 | 0.51 ± 0.091 | 18.1 |
| 2 | 32 | 18 | 0 | 0.49 ± 0.12 | 24.5 |
| 3 | 13 | 9 | 1 | 0.49 ± 0.15 | 30.6 |
| 4 | 26 | 24 | 3 | 0.52 ± 0.12 | 23.6 |
| 5 | 4 | 4 | 2 | 0.50 ± 0.10 | 20.6 |
| 6 | 14 | 10 | 0 | 0.51 ± 0.14 | 27.5 |
| 7 | 2 | 2 | 0 | 0.45 ± 0.037 | 8.4 |
| 8 | 8 | 7 | 0 | 0.54 ± 0.063 | 11.7 |
| 9 | 21 | 14 | 3 | 0.51 ± 0.077 | 15.0 |
| 10 | 34 | 30 | 3 | 0.51 ± 0.051 | 10.0 |
| 11 | 7 | 5 | 2 | 0.55 ± 0.086 | 15.1 |
| 12 | 6 | 3 | 0 | 0.49 ± 0.16 | 32.3 |
| 13 | 15 | 11 | 0 | 0.48 ± 0.070 | 14.5 |
| 14 | 16 | 13 | 2 | 0.52 ± 0.058 | 11.2 |
| 15 | 11 | 11 | 1 | 0.54 ± 0.11 | 20.5 |
| Mean | | | | 0.51 ± 0.026 | 18.9 ± 7.6 |

**Table S3.** Data for aGUVs obtained from probe entry experiments: number of vesicles ($N$), number of phase separated vesicles including modulated ($N_{2\phi}$), number of modulated vesicles ($N_{mod}$), Ld phase area fraction ($f_{Ld}$), relative uncertainty in Ld area fraction ($\sigma_{f_{Ld}}/f_{Ld}$).

| Prep | $N$ | $N_{2\phi}$ | $N_{mod}$ | $f_{Ld}$ | $\sigma_{f_{Ld}}/f_{Ld}$ (%) |
|---|---|---|---|---|---|
| 1 | 24 | 22 | 2 | 0.52 ± 0.051 | 9.9 |
| 2 | 26 | 20 | 1 | 0.56 ± 0.14 | 25.1 |
| 3 | 4 | 3 | 0 | 0.52 ± 0.055 | 10.5 |
| 4 | 5 | 3 | 0 | 0.47 ± 0.083 | 17.8 |
| 5 | 50 | 39 | 7 | 0.52 ± 0.075 | 14.3 |
| 6 | 29 | 27 | 3 | 0.53 ± 0.063 | 11.7 |
| 7 | 3 | 2 | 0 | 0.62 ± 0.0020 | 0.3 |
| 8 | 34 | 29 | 1 | 0.53 ± 0.11 | 21.2 |
| 9 | 3 | 2 | 0 | 0.52 ± 0.12 | 23.3 |
| 10 | 16 | 11 | 0 | 0.49 ± 0.13 | 25.8 |
| 11 | 36 | 26 | 0 | 0.62 ± 0.13 | 21.7 |
| 12 | 17 | 9 | 1 | 0.56 ± 0.086 | 15.4 |
| Mean | | | | 0.54 ± 0.045 | 16.4 ± 7.6 |

**Table S4.** Data for symmetric control GUVs used in probe entry experiments: number of vesicles ($N$), average vesicle intensity ($\bar{F}$), relative uncertainty in intensity ($\sigma_{\bar{F}}/\bar{F}$), average diameter ($d$), Ld phase area fraction ($f_{Ld}$), relative uncertainty in Ld area fraction ($\sigma_{f_{Ld}}/f_{Ld}$).

| Prep | $N$ | $\bar{F}$ | $\sigma_{\bar{F}}/\bar{F}$ (%) | $d$ ($\mu m$) | $f_{Ld}$ | $\sigma_{f_{Ld}}/f_{Ld}$ (%) |
|---|---|---|---|---|---|---|
| 1 | 44 | 1130 ± 279 | 24.6 | 19 ± 7.6 | 0.50 ± 0.12 | 23.1 |
| 2 | 46 | 1080 ± 186 | 17.2 | 19 ± 3.5 | 0.48 ± 0.10 | 21.1 |
| 3 | 47 | 1380 ± 131 | 9.5 | 24 ± 5.7 | 0.54 ± 0.033 | 6.2 |
| 4 | 43 | 1200 ± 120 | 10.4 | 22 ± 7.5 | 0.52 ± 0.027 | 5.2 |
| 5 | 49 | 1060 ± 175 | 16.5 | 19 ± 4.5 | 0.54 ± 0.053 | 9.8 |
| 6 | 45 | 1100 ± 170 | 15.7 | 19 ± 3.7 | 0.54 ± 0.035 | 6.5 |
| 7 | 44 | 1300 ± 250 | 19.2 | 22 ± 6.1 | 0.56 ± 0.039 | 7.1 |
| 8 | 50 | 1200 ± 150 | 12.9 | 21 ± 6.4 | 0.58 ± 0.052 | 9.0 |
| 9 | 44 | 1290 ± 205 | 16.0 | 21 ± 4.7 | 0.54 ± 0.070 | 12.9 |
| 10 | 46 | 1300 ± 150 | 11.6 | 22 ± 4.5 | 0.55 ± 0.086 | 8.6 |
| 11 | 45 | 1390 ± 135 | 9.8 | 21 ± 6.2 | 0.54 ± 0.032 | 6.0 |
| 12 | 47 | 1400 ± 170 | 12.0 | 21 ± 8.9 | 0.55 ± 0.045 | 8.2 |
| Mean | 46 | 1240 ± 124 | 14.6 ± 0.045% | 21 ± 1.6 | 0.54±0.026 | 10.3 ± 5.9 |

**S1. Deriving the uncertainty in exchange fraction calculated from fluorescence intensity.**

We calculate the fraction of exchanged outer leaflet lipids, $\varepsilon_{obs}$, of an aGUV as follows:

$$\varepsilon_{exit} = 2\left(1 - \frac{F_A}{\bar{F}_S}\right), \qquad S1a$$

$$\varepsilon_{entry} = 2\frac{F_A}{\bar{F}_S}, \qquad S1b$$

where Eq. S1a or S1b are appropriate for a probe exit or probe entry experiment, respectively. In Eqs. S1, $F_A$ is the total fluorescence intensity of an individual aGUV and $\bar{F}_S$ is the average fluorescence intensity of $N_S$ symmetric control GUVs at a probe concentration representing complete exchange of both leaflets:

$$\bar{F}_S = \frac{1}{N_S}\sum_{i=1}^{N_S} F_{S,i}\,. \qquad S2$$

$\bar{F}_S$ is thus a reference intensity for calculating outer leaflet exchange; for either mode of probe transfer, the intensity of an asymmetric GUV at 100% outer leaflet exchange is half the value of $\bar{F}_S$.

Defining $f = F_A/\bar{F}_S$, we rewrite Eqs. S1:

$$\varepsilon_{exit} = 2 - 2f, \qquad S3a$$

$$\varepsilon_{entry} = 2f. \qquad S3b$$

Using standard rules for error propagation, the uncertainty in $f$ is:

$$\sigma_f = f\sqrt{\left(\frac{\sigma_{F_A}}{F_A}\right)^2 + \left(\frac{\sigma_{\bar{F}_S}}{\bar{F}_S}\right)^2}\,. \qquad S4$$

Rearranging Eqs. 3 to solve for $f$, we have:

$$f_{exit} = 1 - \varepsilon_{exit}/2\,, \qquad S5a$$

$$f_{entry} = \varepsilon_{entry}/2\,. \qquad S5b$$

Inserting this result into Eq. S4 gives:

$$\sigma_{f_{exit}} = (1 - \varepsilon_{exit}/2) \sqrt{\left(\frac{\sigma_{F_A}}{F_A}\right)^2 + \left(\frac{\sigma_{\bar{F}_S}}{\bar{F}_S}\right)^2}, \qquad S6a$$

$$\sigma_{f_{entry}} = \varepsilon_{entry}/2 \sqrt{\left(\frac{\sigma_{F_A}}{F_A}\right)^2 + \left(\frac{\sigma_{\bar{F}_S}}{\bar{F}_S}\right)^2}. \qquad S6b$$

Applying the rules for error propagation to Eqs. 3, we have:

$$\sigma_{\varepsilon_{exit}} = 2\sigma_{f_{exit}} = (2 - \varepsilon_{exit}) \sqrt{\left(\frac{\sigma_{F_A}}{F_A}\right)^2 + \left(\frac{\sigma_{\bar{F}_S}}{\bar{F}_S}\right)^2}, \qquad S7a$$

$$\sigma_{\varepsilon_{entry}} = 2\sigma_{f_{entry}} = \varepsilon_{entry} \sqrt{\left(\frac{\sigma_{F_A}}{F_A}\right)^2 + \left(\frac{\sigma_{\bar{F}_S}}{\bar{F}_S}\right)^2}. \qquad S7b$$

The expression under the radical symbol represents a propagation of the relative error inherent to measuring the total fluorescence intensity of a single aGUV (the first term) with the uncertainty in the average intensity of a *population* of symmetric GUVs (the second term). In other words, the magnitude of the second term can be made arbitrarily small if enough sGUVs are measured. In our experiments, we typically measured 50 symmetric vesicles to determine $\bar{F}_S$. The average relative uncertainty in fluorescence of an individual GUV, $\sigma_{F_S}/F_S$, was about 15% (Tables S1 and S4). The relative uncertainty in the *mean* intensity of 50 GUVs is then:

$$\frac{\sigma_{\bar{F}_S}}{\bar{F}_S} = \frac{1}{\sqrt{N_S}} \frac{\sigma_{F_S}}{F_S} \approx \frac{0.15}{\sqrt{50}} = 0.021.$$

If we assume that the relative uncertainty in the total intensity of an individual aGUV is similar to that of an individual sGUV (i.e., $\sigma_{F_A}/F_A \approx \sigma_{F_S}/F_S = 0.15$), then the expression under the radical in Eqs. S7 is:

$$\sqrt{\left(\frac{\sigma_{F_A}}{F_A}\right)^2 + \left(\frac{\sigma_{\bar{F}_S}}{\bar{F}_S}\right)^2} \approx \sqrt{0.15^2 + 0.021^2} = 0.151,$$

i.e., the propagated error is dominated by the first term in the radical. We can then simplify Eqs. S7:

$$\sigma_{\varepsilon_{exit}} \approx (2 - \varepsilon_{exit}) \frac{\sigma_{F_A}}{F_A}, \qquad S8a$$

$$\sigma_{\varepsilon_{entry}} \approx \varepsilon_{entry} \frac{\sigma_{F_A}}{F_A}. \qquad S8b$$
